# Stress granule dysfunction via chromophore-associated light inactivation

**DOI:** 10.1101/2023.08.12.553066

**Authors:** Takumi Koizumi, Ai Fujimoto, Haruka Kawaguchi, Tsumugi Kurosaki, Akira Kitamura

**Affiliations:** Laboratory of Cellular and Molecular Sciences, Graduate School of Life Science, Hokkaido University, Sapporo 001-0021, Japan; Laboratory of Cellular and Molecular Sciences, Faculty of Advanced Life Science, Hokkaido University, Sapporo 001-0021, Japan; PRIME, Japan Agency for Medical Research and Development, Chiyoda-ku, Tokyo 100-0004, Japan

**Keywords:** stress granule, G3BP1, TDP-43, cell viability, chromophore-associated light inactivation

## Abstract

Stress granules (SGs) are cytoplasmic condensates composed of various proteins and RNAs that protect translation-associated machinery from harmful conditions during stress. However, the method of spatio-temporal inactivation of condensates such as SGs in live cells to study cellular phenotypes is still in the process of being demonstrated. Here, we show that the inactivation of SG by chromophore-associated light inactivation (CALI) using a genetically encoded red fluorescence protein (SuperNova-Red) as a photosensitizer leads to differences in cell viability during recovery from hyperosmotic stress. CALI delayed the disassembly kinetics of SGs during recovery from hyperosmotic stress. Consequently, CALI could inactivate the SGs, and the cellular fate due to SGs could be analyzed. Furthermore, CALI is an effective spatiotemporal knockdown method for intracellular condensates/aggregates and would contribute to the elucidation of importance of such condensates/aggregates.

## 1 Introduction

Stress granules (SGs) are membraneless organelles; they are biomolecular condensates composed of proteins and RNAs that emerge in the cytoplasm under cellular stresses such as hyperosmotic, heat, and oxidative stress ^1, 2^. After being released from stress, SGs are reversibly disassembled ^1^.

Although the function of SGs remains elusive, they have long been proposed to play a role in protecting translation-associated RNAs from harmful conditions under stress because translation-associated proteins and RNAs, such as ribosomes, pre-translation complexes with mRNA, and various translation-associated RNA-binding proteins (RNPs), are widely sequestered into SGs ^3-5^.

Therefore, SGs are believed to play a cytoprotective role under stress conditions. SGs contain a biphasic structure with a stable “core” phase surrounded by a less concentrated “shell” phase ^6^. Ras-GTPase-activating protein SH3 domain-binding protein 1 (G3BP1) is an RNA-binding protein essential for SG formation, which can primarily be found in the core phase ^1, 2^. Loss of G3BP1 increases the number of ubiquitinated proteins and toxic aggregates ^7, 8^ and inhibits the proliferation of cancer cells ^9^. Transactivation response DNA/RNA-binding protein 43 kDa (TDP-43) is an amyotrophic lateral sclerosis-associated aggregation-prone protein and a component of SGs. One starting point for the pathological aggregation of TDP-43 is its localization to and interaction with SG components ^10^. The genetic depletion of endogenous G3BP1 reduces SG formation efficiency ^11^. However, it is still unknown whether the inactivation of SG proteins after formation of SGs causes severe defects in SGs. To elucidate the mechanisms underlying this issue, the spatiotemporal inactivation of the SGs, rather than the genetic knockout/knockdown of genes of interest, is required because SGs may no longer be formed by conventional genetic depletion methods.

Chromophore-assisted light inactivation (CALI) is used for the spatiotemporal inactivation of specific compartments of live cells ^12, 13^. Absorption of light by the chromophore in phototoxic photosensitizers generates reactive oxygen species (ROS), which, in turn, inactivates photosensitizer-proximate biomolecules. Although various photosensitizers have been used, fluorescent protein-based photosensitizers can be labeled at a defined molar ratio (e.g., 1:1) with the target protein, expressed in cells via gene transfection, and localized to specific subcellular compartments ^14^.

SuperNova-Red (SNR) is a monomeric photosensitizer and a red fluorescent protein with ROS production ^15^. Therefore, we hypothesized that the spatiotemporal inactivation of SGs can be achieved by tagging the SNR to specific proteins within specific compartments in SGs under stress.

In this report, we show how CALI changes the cell viability and kinetics of SG disassembly using SNR to G3BP1 and TDP-43 in SGs as targets during the recovery process after hyperosmotic stress.

## 2 Results

### 2.1 After CALI, SG-containing cells showed low viability during recovery from hyperosmotic stress

To demonstrate whether SGs play a cytoprotective role, we established a procedure to observe the viability of murine neuroblastoma Neuro-2a (N2a) cells expressing a protein target in CALI under the control of irradiation time and intensity during the recovery period after hyperosmotic stress caused by 290 mM NaCl (Figure S1). Under this condition, cytoplasmic SG formation is induced through macromolecular crowding by compressed cell volume followed by phase separation and changes in cytoplasmic rheology ^16-18^. First, we tested whether the SNR monomers induce cell death in the recovery medium after hyperosmotic stress. After light irradiation, no SNR fluorescence was detected, indicating that SNR was efficiently photobleached (Figure S2A). Photobleaching indicates that the amount of ROS production from fluorophores has been saturated because ROS is generated upon absorption of photons. ^19^. The viability of cells expressing SNR monomers decreased to 86% at 600 min after the start of the time-lapse observation (Figure S2B), indicating that neither ROS from SNR monomers nor the cellular recovery process from hyperosmotic stress caused cell death.

Next, as G3BP1 localizes in the core phase of SGs, we chose G3BP1 as a target for the inactivation of SGs in CALI. Cells with and without SGs of G3BP1-SNR were observed when cells were exposed to hyperosmotic stress (Figure 1A). This is due to using a mild hyperosmotic stress condition. In this condition, G3BP1-SNR was not overexpressed in N2a cells (Figure S3). Hence, we compared the proportion of surviving cells harboring SGs with that of cells not harboring SGs during recovery from hyperosmotic stress. As negative controls, almost all cells without light irradiation were alive regardless of G3BP1-SNR SG formation (green and orange lines, Figure 1B), indicating that neither the recovery process from hyperosmotic stress nor time-lapse observation affected the survival of the cells expressing G3BP1-SNR. In contrast, cells exposed to hyperosmotic stress with light irradiation gradually died in both SG-positive and SG-negative cell groups during recovery from hyperosmotic stress. The rate of cell death up to 200 min is higher in SG-negative cells than SG-positive cells, though some SG-negative cells seemed to survive (blue and magenta lines, Figure 1B). Therefore, we successfully confirmed the reduction in cell viability induced by CALI in cells expressing G3BP1-SNR, and that SG-positive cells were more likely to be affected by CALI-induced cell death than SG-negative cells.

**Figure 1.**
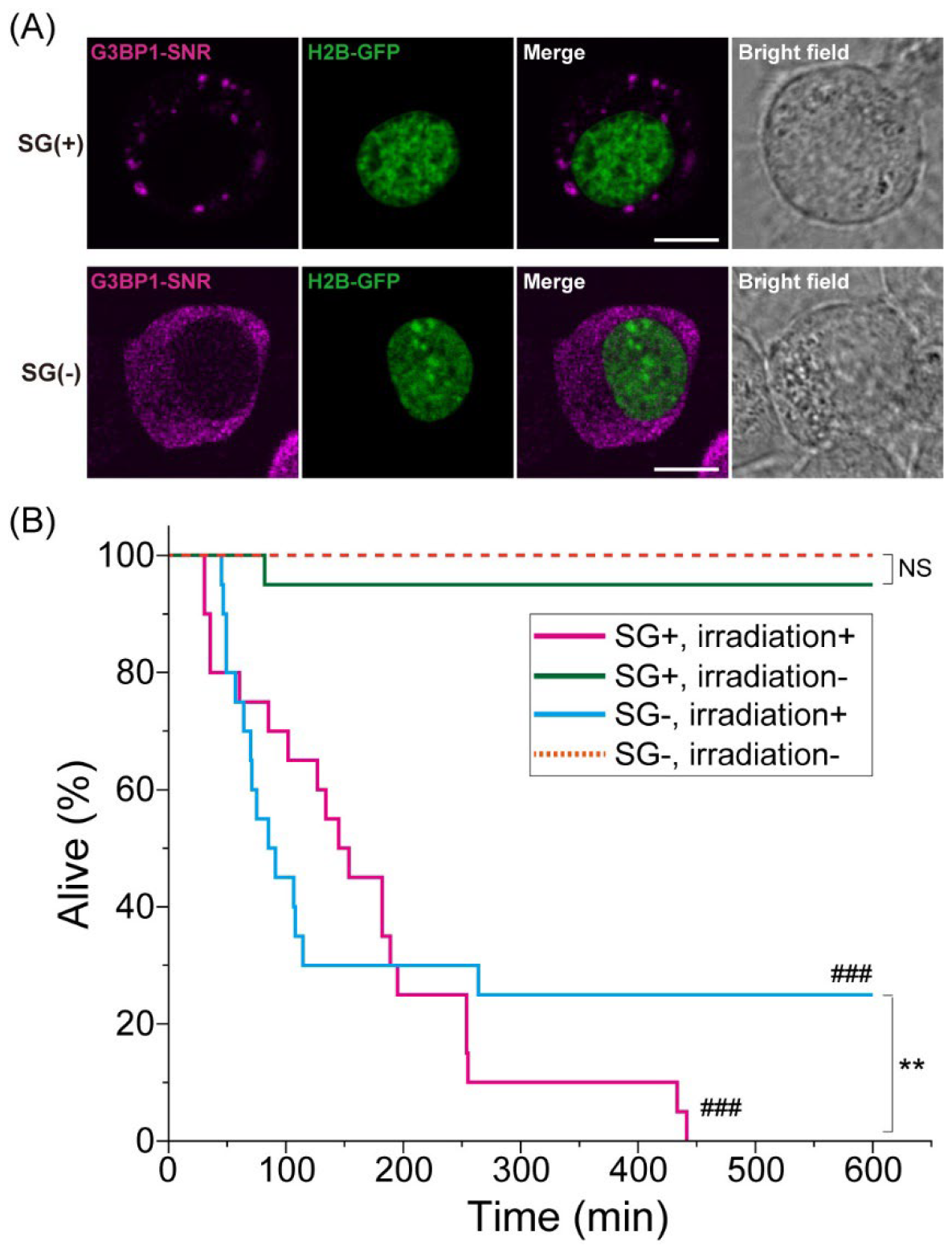
Cell viability after chromophore-associated light inactivation (CALI) targeting G3BP1 in Neuro-2a cells recovering from hyperosmotic stress. (A) Fluorescence images of SG-positive and SG-negative cells expressing SuperNova-Red (SNR)-tagged G3BP1 (G3BP1-SNR; magenta) and Histon H2B-GFP (green) before light irradiation for CALI. (B) Time-course cell viability plot (600 min) in the recovery medium after hyperosmotic stress with or without light irradiation for CALI (20 cells). *P*-values were obtained using the generalized Wilcoxon test (Gehan-Breslow method). ###*p* < 0.001 (compared to the group without light irradiation); ^**^*p* < 0.01 between the lines; NS, not significant (*p* ≥ 0.05).

As TDP-43 localizes in the SGs ^10^, we chose it as a target for the inactivation of SGs in CALI. We also tracked cell viability after light irradiation using the same procedure as G3BP1-SNR, where N2a cells with and without TDP43-SNR SGs were observed (Figure 2A). In this condition, TDP43-SNR was not overexpressed in N2a cells (Figure S3). Similar to G3BP1-SNR-expressing cells, cell death was not observed in TDP43-SNR-expressing cells, regardless of SGs, when no light was used as a negative control (green and orange lines, Figure 2B). In contrast, significant cell death was observed in cells harboring SGs of TDP43-SNR; most cells that did not harbor TDP43-SNR SGs remained alive (blue and magenta lines, Figure 2B). Furthermore, a higher proportion of CALI-induced cell death was observed in cells expressing G3BP1-SNR than in those expressing TDP43-SNR (Figure S4).

**Figure 2.**
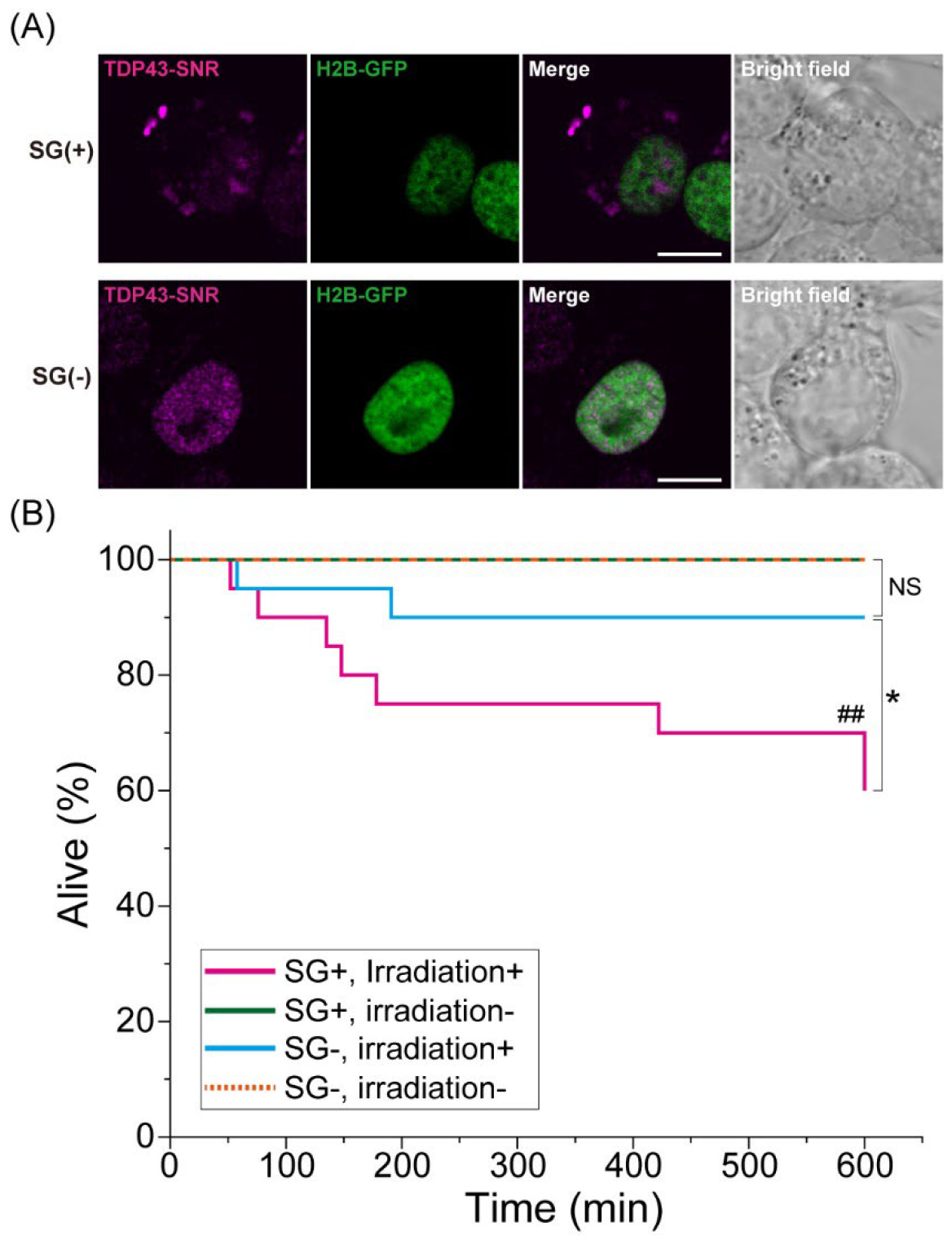
Cell viability after chromophore-associated light inactivation (CALI) targeting TDP-43 in Neuro-2a cells recovering from hyperosmotic stress. (A) Fluorescence images of SG-positive and SG-negative cells expressing SuperNova-Red (SNR)-tagged TDP-43 (TDP43-SNR; magenta) and Histon H2B-GFP (green) before light irradiation for CALI. (B) Time-course cell viability plot (600 min) in the recovery medium after hyperosmotic stress with or without light irradiation for CALI (20 cells). *P*-values were obtained using the generalized Wilcoxon test (Gehan-Breslow method). ##*p* < 0.01 (compared to the group without light irradiation); ^*^*p* < 0.05 between the lines; NS, not significant (*p* ≥ 0.05).

Furthermore, to detect ROS production after CALI for SNR in live cells, a cell-permeable and fluorescent ROS detection reagent was used. After introducing the ROS detection reagent into the cells, the green fluorescence intensity of the ROS detection reagent increased in G3BP1/TDP43-SNR and SNR-expressing cells with the fluorescence loss of SNR after light irradiation for CALI, whereas that of the ROS detection reagent did not increase in cells without expression of SNRs (Figure S5).

This result suggests that SNR generates ROS by light irradiation in live cells.

### 2.2 CALI inhibits SG disassembly

As SGs are gradually disassembled during recovery from hyperosmotic stress ^20^, we investigated whether CALI-induced G3BP1 or TDP-43 inactivation in SGs affected the disassembly of SGs. To observe the subcellular localization of G3BP1 or TDP-43 after the photobleaching of SNR for CALI, N2a cells co-expressing meGFP-tagged and SNR-tagged G3BP1/TDP-43 were subjected to hyperosmotic stress to form SGs in the cytoplasm, and then irradiated with light immediately after replacement with the recovery medium. Immediately after light irradiation in CALI, the SNR fluorescence completely disappeared; however, the subcellular localization of GFP-tagged G3BP1 and TDP-43 was not affected (Figure S6). Next, the disassembly of G3BP1- and TDP-43-containing SGs was observed using time-lapse fluorescence microscopy (Figure 3A). Although G3BP1-containing SGs disappeared at 30 min in the cell without light irradiation, they remained in the light-irradiated cell (Figure 3A). In contrast, TDP-43-containing SGs remained in the cells for a long time compared to G3BP1-containing SGs (Figure 3A). To compare the difference quantitatively, we plotted the time-dependent change of SG number in the cells (Figures 3, B & C). G3BP1-containing SGs were gradually disassembled with and without light irradiation in the recovery medium.

**Figure 3.**
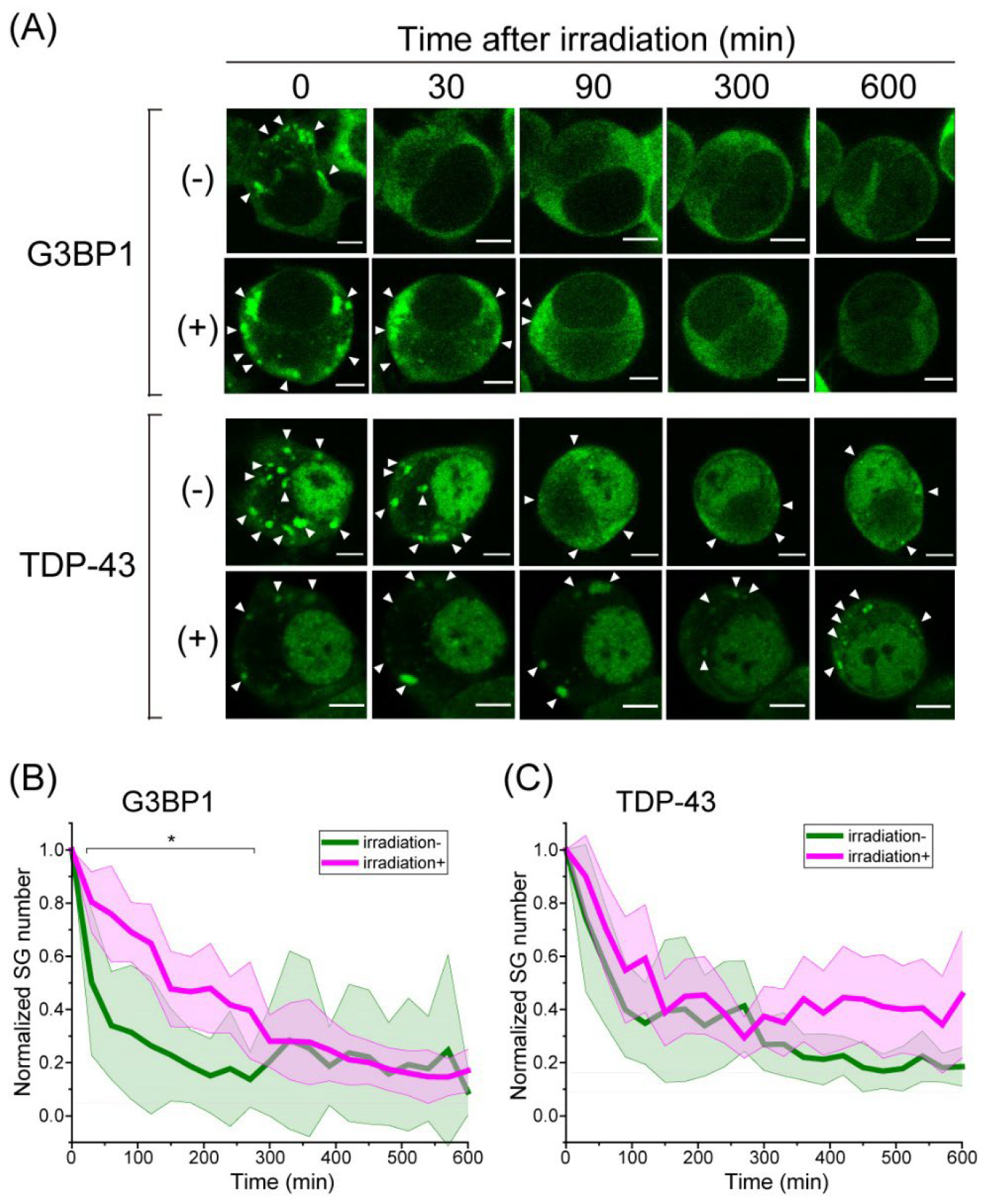
Disassembly of SGs containing G3BP1-GFP or TDP43-GFP after chromophore-associated light inactivation (CALI) targeting G3BP1-SNR or TDP43-SNR, respectively. (A) Time-lapse observation of Neuro-2a cells harboring SGs of G3BP1-GFP or TDP43-GFP with (+) or without (-) light irradiation. Bar = 5 μm. White arrow heads indicate the position of SGs. (B) Plot showing the decrease in the normalized number of SGs containing G3BP1-GFP with (magenta) or without (green) light irradiation (mean ± 95%CI; 15 cells). The number of SGs containing G3BP1-GFP was normalized by that immediately after the start of observation (the mean number of SGs in a cell was 12 for irradiation (-) and 16 for irradiation (+)), and the mean values of decreasing ratio at each observation time are shown. *P*-values were obtained using Student’s t-test. ^*^*p* < 0.05 at the time indicated on the lines. (C) Plot showing the decrease in the normalized number of SGs containing TDP43-GFP with (magenta) or without (green) light irradiation (mean ± 95%CI; 15 cells). The number of SGs containing TDP43-GFP was normalized by that immediately after the start of observation (the mean number of SGs in a cell was 14 for irradiation (-) and 12 for irradiation (+)), and the mean values of decreasing ratio at each observation time are shown. *P*-values were obtained using Student’s t-test.

However, in the light-irradiated cells, SG disassembly speed was slightly delayed up to ∼270 min in the recovery medium, and the population of cells harboring SGs decreased to the same extent with and without light irradiation (>300 min) (Figure 3B). The TDP-43-containing SGs with and without light irradiation gradually disassembled at a similar speed in the recovery medium (Figure 3C). In summary, CALI slightly inhibited disassembly of G3BP1-containing SGs but did not that of TDP-43-containing ones.

## 3 Discussion

Here, we showed that CALI using SNR to target G3BP1 and TDP-43 in SGs increased cell death (Figures 1&2) and inhibited SG disassembly by G3BP1 inactivation (Figure 3). First, we determined the possible mechanisms by which CALI inhibits SG disassembly. Light-irradiated SNR generates both superoxide and singlet oxygen as ROS; however, the quantum yield of SNR to generate singlet oxygen is approximately 15-fold higher than that to generate superoxide ^19^. The higher efficiency of SNR in generating singlet oxygen compared to superoxide is similar to other red fluorescent proteins, such as mCherry ^21^ and KillerRed ^22^ but the highest among these proteins ^15, 19^. The lifetime of singlet oxygen in water was reported at 3.7 μs ^23^. However, the travel distance of singlet oxygen in cells is believed to be less than 2 nm because singlet oxygen reacts with water and other biomolecules ^24^; hence, singlet oxygen can spread only within the local environment ^25^. Because the short diameter of the fluorescent protein-conserved β-barrel of SNR is approximately 1.3 nm ^15^, the singlet oxygen generated from SNR is consumed by biomolecules close to the outside of the β-barrel.

Singlet oxygen damages proteins, peptides, and nucleic acids in various ways ^26-30^. In proteins and peptides, singlet oxygen can react with the side chains of amino acids (e.g., Trp, Tyr, His, Met, and Cys), resulting in the chemical modification of these amino acids, backbone fragmentation, cross-linking, and subsequent aggregation ^26, 31^. In nucleic acids, guanosine is a major oxidization target of singlet oxygen ^29, 30^. Because SGs are densely packed with various proteins and RNAs ^1, 2^, singlet oxygen generated from G3BP1-SNR or TDP43-SNR may act on their proximate proteins and RNAs in SGs, leading to side chain modifications, backbone fragmentations, and cross-linking of SG components in the phase where SNR-tagged proteins are localized (Figure 4). These effects do not produce an immediate change or instantaneous disappearance of SGs (Figure S6). Since G3BP1 is a core protein in SGs, its inactivation may inhibit SG disassembly (Figure 3). In contrast, since TDP-43 is not essential to SG formation, its inactivation may not inhibit SG disassembly (Figure 3). As previously reported, the most easily interpreted cause of difficulty in SG disassembly is intermolecular cross-linking ^32-34^. Another possibility is that molecular chaperones such as HSP70 are incorporated in SGs to protect and possibly repair proteins that unfold during various types of stress and are required for the disassembly of SGs ^3^. Inactivation of such molecular chaperones by CALI would cause defects in SG disassembly. As the abundance, conformations, modifications, and interactions of RNA may control the assembly and disassembly of SGs ^1, 35^, singlet oxygen-induced guanosine oxygenation in RNA may retard SG disassembly process.

**Figure 4.**
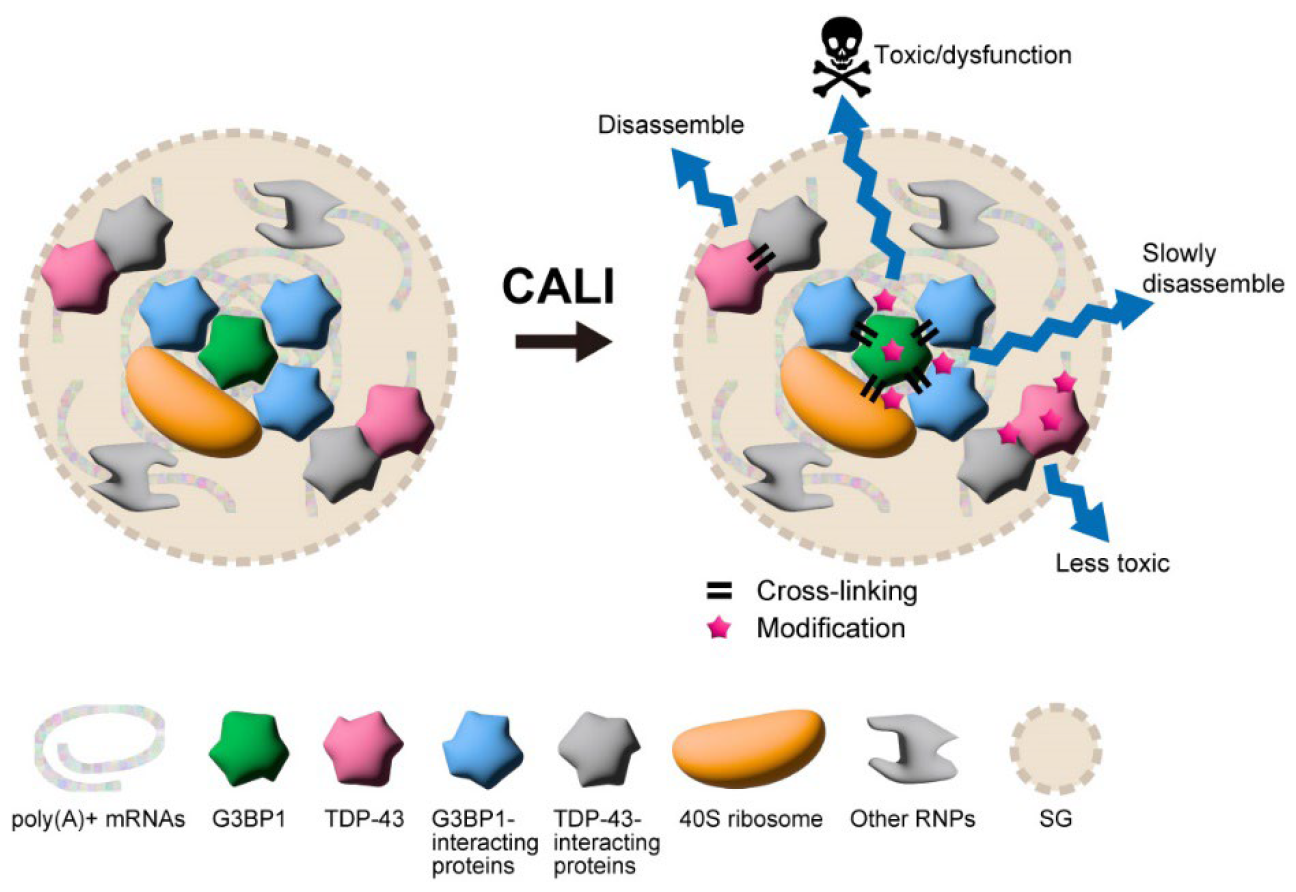
The possible mechanism of internal damage in SGs caused by light irradiation.

Because cross-linking is a covalent bond, the dissociation of biomolecules is less likely to occur.

Interestingly, the disassembly kinetics differed between G3BP1 and TDP-43 (Figure 3). Although G3BP1 localizes to the “core” phases in SG and its function against SGs is essential, sub-localization and essential effects of TDP-43 in the SGs are uncharacterized. Proteins and RNAs cross-linked from TDP-43 may have relatively little effect on SG maintenance and cell survival. In contrast, when SG itself collapses by CALI for G3BP1, it is likely to dissociate less if it is cross-linked (Figure 4).

However, light-irradiated G3BP1-harboring SGs disassembled even after a delay, so there may be a repair system for cross-linked species during SG disassembly. Although singlet oxygen was artificially generated from SNR in this experiment, singlet oxygen and other ROS can be generated by intracellular redox reactions in physiological environments ^36^. Singlet oxygen and other ROS generated in physiological states form intermolecular cross-links in SGs. In other words, this study attempted to mimic the effects of singlet oxygen-promoted inactivation of intracellular structures such as SGs.

Next, we discuss the influence of CALI outside SGs (i.e., in the cytoplasm and nucleus). The G3BP1-SNR and TDP-43-SNR used in this study were localized not only in SGs but also in the cytoplasm and nucleus. Because whole cells were irradiated with light, ROS may have been generated in the cytoplasm and nucleus. Our results showed that cell death was more pronounced after CALI in SG-positive cells for both G3BP1-SNR and TDP-43-SNR (Figures 1&2), suggesting that SG inactivation was more strongly associated with cell death than defects in these proteins localized in the cytoplasm or nucleus. However, CALI in cells not harboring SGs of G3BP1-SNR also induced cell death (Figure 1B), suggesting that the inactivation of G3BP1 in the cytoplasm and nucleus may be involved in cell death. Loss of G3BP1 increases protein ubiquitination and the aggregation of toxic huntingtin ^7, 8^ and diminishes the proliferation of breast cancer cells ^9^; the inactivation of G3BP1 by CALI likely leads to proteostasis dysregulation and eventual cell death. In other words, G3BP1 plays a cytoprotective role in promoting cell viability. Therefore, cell death may appear gradually with the progress of SG disassembly, and eventually, all of them may be dead. Loss of TDP-43 leads to various cellular dysfunctions, such as the misregulation of RNA splicing and its quality control, mitochondrial dysfunction, and cell death ^37-40^. However, cell death during TDP-43 knockdown was observed at approximately 120 h after siRNA introduction ^40^. This suggests that cell death by TDP-43 depletion may not be an acute response but rather via the accumulation of various dysfunctions. CALI of TDP43-SNR in cells not harboring SGs did not result in a dramatic increase in cell death (Figure 2B), consistent with the slow rate of cell death observed in TDP43-depleted cells. This supports the specificity of CALI for the protein of interest and its localization.

Finally, we discuss a possible mechanisms of cell death induced by CALI in SGs. SGs attenuate apoptotic cell death via p53 inhibition and the recruitment of ROCK1 and RACK1 to SGs ^41-43^. SG formation regulates several canonical signaling pathways (such as NF-κB, mTORC1, and PKR) and pro-survival functions during the acute stress response ^1, 4^. The dysfunction of SGs induced by CALI may reactivate transiently inhibited apoptotic signaling pathways. The CALI-induced aberrant RNA in SGs may disturb cellular functions. mRNA translation is stalled as a pre-initiation complex in SGs, as translation is traditionally considered silent during SG formation ^1^. Although some mRNAs are translationally silent, a host of RNA molecules undergo translation in SGs ^44^. Therefore, the translation dysfunction required for stress recovery may be delayed by aberrant RNA induced by CALI in SGs, leading to cell death.

Consequently, here, we showed that CALI of two SG components (G3BP1 and TDP-43) using SNR led to cell death. Our results suggest that CALI can inactivate various intracellular condensates composed of proteins and RNA in a spatiotemporally controlled manner, and the subsequent cellular fate can be analyzed. Furthermore, we believe that CALI is an effective method that contributes to identifying intercellular condensates/aggregates as hubs controlling signal transduction. However, the molecular mechanism of inactivation by CALI remains unknown, and its identification remains an important open question.

## 4 Methods

### Plasmids

To create the expression plasmids of proteins that SNR was tagged at the C-terminus of G3BP1 and TDP-43, SNR cDNA fragment (gift from Prof. T. Nagai) was inserted into the *Bam*HI and *Not*I sites of pcDNA3.1(+) (Thermo Fisher Scientific, Waltham, MA, USA) (pcDNA-SNR). Next, the DNA fragments of human TDP-43 obtained from the TDP-43 expression plasmid as previously reported ^45^ and G3BP1 from Addgene (Clone#129339, Watertown, MA, USA) were inserted into the *Hind*III and *Bam*HI sites of pcDNA-SNR (pTDP43-SNR and pG3BP1-SNR, respectively). To create monomeric enhanced green fluorescent protein (meGFP)-tagged TDP-43 or G3BP1 expression vectors, the DNA fragment of the SNR of pTDP43-SNR and pG3BP1-SNR was cut and substituted with that of meGFP digested from pmeGFP-N1 ^45^ using the *Bam*HI and *Not*I sites (pTDP43-GFP and pG3BP1-GFP, respectively). The expression plasmid for meGFP-tagged histone H2B (pBOS-H2B-GFP) and pCAGGS were the same as that used previously ^45, 46^.

### Cell Culture and Transfection

Neuro-2a murine neuroblastoma (N2a) cells were obtained from the American Type Culture Collection (ATCC; Manassas, VA, USA) and maintained in Dulbecco’s Modified Eagle Medium (DMEM) (D5796, Sigma-Aldrich, St. Louis, MO, USA) supplemented with 10% heat-inactivated fetal bovine serum (FBS) (12676029, Thermo Fisher Scientific), 100 U/mL penicillin G (Sigma-Aldrich), and 0.1 mg/mL streptomycin (Sigma-Aldrich), as previously described^45^. One day before transfection, 2.0 × 10^5^ N2a cells were transferred to a glass bottom dish (3910-035, IWAKI-AGC Technoglass, Shizuoka, Japan). The mixture of expression plasmids was transfected using Lipofectamine 2000 (Thermo Fisher Scientific) according to the manufacturer’s protocol. Details on the amounts of the plasmids are described in Table S1. After 24 h of incubation, the spent medium was replaced with Opti-MEM I (Thermo Fisher Scientific) or Opti-MEM I containing 290 mM NaCl (hyperosmotic medium).

### Western blotting

N2a cells were lysed in a PBS containing 1% SDS, 0.25 U/μL benzonase, and 1× protease inhibitor cocktail (Sigma-Aldrich). After the centrifugation (20,400 g, 10 min., 4°C), the supernatants were recovered. SDS-PAGE sample buffer-mixed lysates were denatured at 98°C for 5 min. Samples were applied to a 12.5% polyacrylamide gel (ATTO, Tokyo, Japan) and subjected to electrophoresis in an SDS-containing buffer. The proteins were transferred to a PVDF membrane (Cytiva, Marlborough, MA). The membrane was incubated in 5% skim milk in PBS-T for background blocking for 1 h. After incubation with an anti-G3BP1 antibody (#ab181150, Abcam, Cambridge, UK), an anti-TDP-43 antibody (#3448, Cell Signaling Technology, Danvers, MA, USA), or an anti-KillerRed antibody (#AB961, Evrogen, Moscow, Russia) in CanGet signal immunoreaction enhancer solution 1 (TOYOBO, Osaka, Japan), horseradish peroxidase-conjugated secondary antibody that is appropriate for primary antibodies (#111-035-144; Jackson Immuno Research, West Grove, PA, USA) was inclubated in 5% skim milk in PBS-T. As a loading control, horseradish peroxidase-conjugated anti-α-tubulin antibody (#HRP-66031, Proteintech, Rosemont, IL, USA) was incubated in 5% skim milk in PBS-T. Chemiluminescence signals were acquired using a ChemiDoc MP imager (Bio-Rad, Hercules, CA, USA).

### Stress granule formation, light irradiation for CALI, and time-lapse observation

To generate SGs under hyperosmotic stress, N2a cells were incubated in a hyperosmotic medium for 3 h. After the hyperosmotic medium was removed, the cells were washed thrice in Hank’s balanced salt solution, and then the medium was exchanged with Opti-MEM I (recovery medium) containing 0.3 mM DRAQ7 (BioStatus, Leicestershire, UK) as an indicator for dead cells. After medium exchange, the glass-based dish was immediately placed on a microscope stage within 2 min. Light irradiation and time-lapse observations were performed using an Axioobserver Z1 inverted microscope (Carl Zeiss, Jena, Germany) equipped with an LSM510 META laser scanning confocal unit (Carl Zeiss), a C-Apochromat 40×/1.2NA W Korr. UV-VIS-IR water immersion objective (Carl Zeiss), laser sources, a mercury lamp, and a heat stage incubator (at 37°C supplemented with 5% CO_2_). The field stop was set to a minimum. For CALI, cells 1–5 expressing SNR-tagged proteins were simultaneously irradiated with light from a mercury lamp through a 540–580 nm bandpass filter (ET560/40x; Chroma Technology Corporation, Bellows Falls, VT, USA) and a 585 nm dichroic mirror (T585lpxr; Chroma Technology Corporation) for 2 min. The aperture diaphragm was adjusted so that the light intensity was 0.9 mW just above the objective lens. After the CALI of SNR, bright-field and DRAQ7 fluorescence images of the irradiated cells were acquired at 30-minute intervals for up to 600 min (Figure S1). The DRAQ7-stained cells were excited at 633 nm. The excitation and emission beams were split using a beam splitter (HFT405/488/543/633). Fluorescence was detected using photomultiplier tubes through a 650 nm long-pass filter (LP650). The diameter of the pinhole for DRAQ7 was 92 μm. The image size was 1024 × 1024 pixels and the zoom factor was 3. DRAQ7-positive cells were considered dead. To observe SG disassembly, GFP was excited at 488 nm. The excitation and emission beams were split using a beam splitter (HFT488/594). Fluorescence was detected using photomultiplier tubes through a 505–570 nm bandpass filter (BP505–570). The pinhole diameter was 72 μm, the image size was 1024 × 1024 pixels, and the zoom factor was 4.

After light irradiation for CALI, bright-field and GFP fluorescence images of the irradiated cells were acquired at 30-minute intervals for up to 600 min. The number of SGs containing G3BP1-GFP and TDP-43-GFP in a cell was normalized by that immediately after the start of observation, and the values of decreasing ratio at each observation time were calculated, and then the mean of the normalized values of the ratio for observed cells was obtained.

### Confocal microscopy

Image acquisition was performed using an inverted fluorescence microscope, Axioobserver Z1 (Carl Zeiss, Jena, Germany), combined with an LSM 510META system (Carl Zeiss), a C-Apochromat 40×/1.2NA W Korr. UV-VIS-IR water immersion objective (Carl Zeiss), laser sources, a mercury lamp, and a heat stage incubator (at 37°C supplemented with 5% CO_2_). GFP and SNR were excited at 488 nm and 594 nm, respectively. The excitation and emission beams were split using a beam splitter (NFT488/594). Fluorescence was detected using photomultiplier tubes through a 615 nm long-pass filter (LP615). The diameter of the pinhole was 72 μm for GFP and 208 μm for SNR.

### Immunofluorescence microscopy

N2a cells were incubated in a hyperosmotic medium for 3 h. After the hyperosmotic medium was removed, 4% paraformaldehyde buffered with 100 mM Hepes-KOH (pH 7.5) was added for fixation for 30 min at 37°C. The fixed cells were washed thrice in Tris-buffered saline, and then the cells were permeabilized in 0.5% Triton X-100 (Fujifilm Wako Chemicals, Osaka, Japan) buffered with phosphate buffer saline (PBS) for 5 min at 25°C, and then the cells were blocked in PBS containing 5% normal goat serum (Sigma-Aldrich), 0.02% Triton X-100, and 20% Glycerol for 24 h at 4°C. Primary antibodies for G3BP1 (#ab181150, Abcam) or TDP-43 (#3448, Cell Signaling Technology) were treated in the buffer as same as the blocking process for 1 h at 25°C. After washing the primary antibodies thrice in PBS, Alexa Fluor 488-conjugated secondary antibodies (#R37116, Thermo Fisher Scientific) were treated in the buffer as same as the blocking process for 1 h at 25°C. After washing the secondary antibodies thrice in PBS, cells were mounted in PBS. Image acquisition was performed using an inverted fluorescence microscope, Axioobserver Z1 (Carl Zeiss), combined with an LSM 510META system (Carl Zeiss), a C-Apochromat 40×/1.2NA W Korr. UV-VIS-IR water immersion objective (Carl Zeiss), laser sources, a mercury lamp, and a heat stage incubator (at 37°C supplemented with 5% CO_2_). Alexa Fluor 488 and SNR were excited at 488 nm and 594 nm, respectively. The excitation and emission beams were split using a beam splitter (NFT488/594). Fluorescence was detected using photomultiplier tubes through a 505–570 nm bandpass filter (BP505–570) for ROS detection reagents and a 615 nm longpass filter (LP615) for SNR. The diameter of the pinhole was 72 μm for GFP and 172 μm for SNR.

### ROS detection

To detect ROS in the cell, ROS Assay Kit -Photo-oxidation Resistant DCFH-DA-(R253, Dojindo, Kumamoto, Japan) was used referring to the manufacturer’s protocol. Briefly, One day before transfection, 0.2 × 10^5^ N2a cells were transferred to a 96-well glass bottom dish (5866-096, IWAKI-AGC Technoglass). The mixture of expression plasmids was transfected using Lipofectamine 2000 (Thermo Fisher Scientific) according to the manufacturer’s protocol. Details on the amounts of the plasmids are described in Table S1. After 24 h of incubation, the medium was removed, and the cells were washed twice with HBSS. Photo-oxidation Resistant DCFH-DA was diluted in a 1× Loading Buffer included in the kit (10 μM DCFH-working solution). After removing the HBSS, 10 μM DCFH-working solution was added and the cells were incubated for 30 min. The cells were washed twice with HBSS, and the wells were refilled with HBSS. Light irradiation and fluorescence observations were performed using an Axioobserver Z1 inverted microscope (Carl Zeiss) equipped with an LSM510 META laser scanning confocal unit (Carl Zeiss), a C-Apochromat 40×/1.2NA W Korr. UV-VIS-IR water immersion objective (Carl Zeiss), laser sources, a mercury lamp, and a heat stage incubator (at 37°C supplemented with 5% CO_2_). The field stop was set to a minimum. For CALI, cells expressing SNR-tagged proteins were simultaneously irradiated with light from a mercury lamp through a 540–580 nm bandpass filter (ET560/40x; Chroma Technology Corporation) and a 585 nm dichroic mirror (T585lpxr; Chroma Technology Corporation) for 2 min. The aperture diaphragm was adjusted so that the light intensity was 0.9 mW just above the objective lens. After the light irradiation to the cells, The ROS detection reagents and SNR in the cells were excited at 488 nm and 594 nm, respectively. The excitation and emission beams were split using a beam splitter (NFT488/594). Fluorescence was detected using photomultiplier tubes through a 505– 570 nm bandpass filter (BP505–570) for ROS detection reagents and a 615 nm longpass filter (LP615) for SNR. The diameter of the pinhole was 77 μm for GFP and 174 μm for SNR, respectively.

### Statistical analyses

Student’s t-tests were performed using Microsoft Excel. The generalized Wilcoxon test (Gehan-Breslow method) was performed using the Bell Curve for Excel software (Social Survey Research Information Co., Ltd., Tokyo, Japan).

## 5 Author Contributions

TKo: Data curation, Formal analysis, Investigation, Methodology, Validation, Visualization; AF: Data curation, Formal analysis, Investigation, Methodology, Validation, Funding acquisition, Writing – review & editing; HK: Data curation, Formal analysis, Investigation, Methodology, Validation; TKu: Investigation, Methodology, Validation; AK: Conceptualization, Data curation, Formal analysis, Funding acquisition, Investigation, Methodology, Project administration, Resources, Supervision, Validation, Visualization, Writing – review & editing.

## 6 Supporting Information

Additional experimental details, materials, and methods, including photographs of experimental setup

## 7 Acknowledgments

We would like to thank A. Murata for technical support, M. Oura for technical suggestions, and T. Nagai for generously providing us with a plasmid vector for SuperNova-Red. We would like to thank Editage (www.editage.jp) for English language editing.

