## Supplemental Figures for "Stress granule dysfunction via chromophore-associated light inactivation"

### 1 Supplementary Figures and Tables

#### 1.1 Table S1

| Corresponding figure | Plasmid 1 | Amount (µg) | Plasmid 2 | Amount (µg) | Total amount (µg) |
|---|---|---|---|---|---|
| **Figure 1 & 3** | pG3BP1-SNR | 0.7 | pBOS-H2B-GFP | 0.3 | 1.0 |
| **Figure 2 & 3** | pTDP43-SNR | 0.7 | pBOS-H2B-GFP | 0.3 | 1.0 |
| **Figure 4 A & B** | pG3BP1-SNR | 0.7 | pG3BP1-GFP | 0.3 | 1.0 |
| **Figure 4 A & C** | pTDP43-SNR | 0.7 | pTDP43-GFP | 0.3 | 1.0 |
| **Figure S2** | pcDNA-SNR | 0.7 | pBOS-H2B-GFP | 0.3 | 0.1 |
| **Figure S3** | pG3BP1-SNR | 0.7 | pCAGGS | 0.3 | 0.1 |
| **Figure S3** | pTDP43-SNR | 0.7 | pCAGGS | 0.3 | 0.1 |
| **Figure S4** | pG3BP1-SNR | 0.07 | pCAGGS | 0.03 | 0.1 |
| **Figure S4** | pTDP43-SNR | 0.07 | pCAGGS | 0.03 | 0.1 |
| **Figure S4** | pcDNA-SNR | 0.07 | pCAGGS | 0.03 | 0.1 |

**Table S1.** The amount and combination of plasmids for each experiment per single glass-base dish. See the Materials & Methods section for details on each plasmid.

#### 1.2 Supplementary Figures

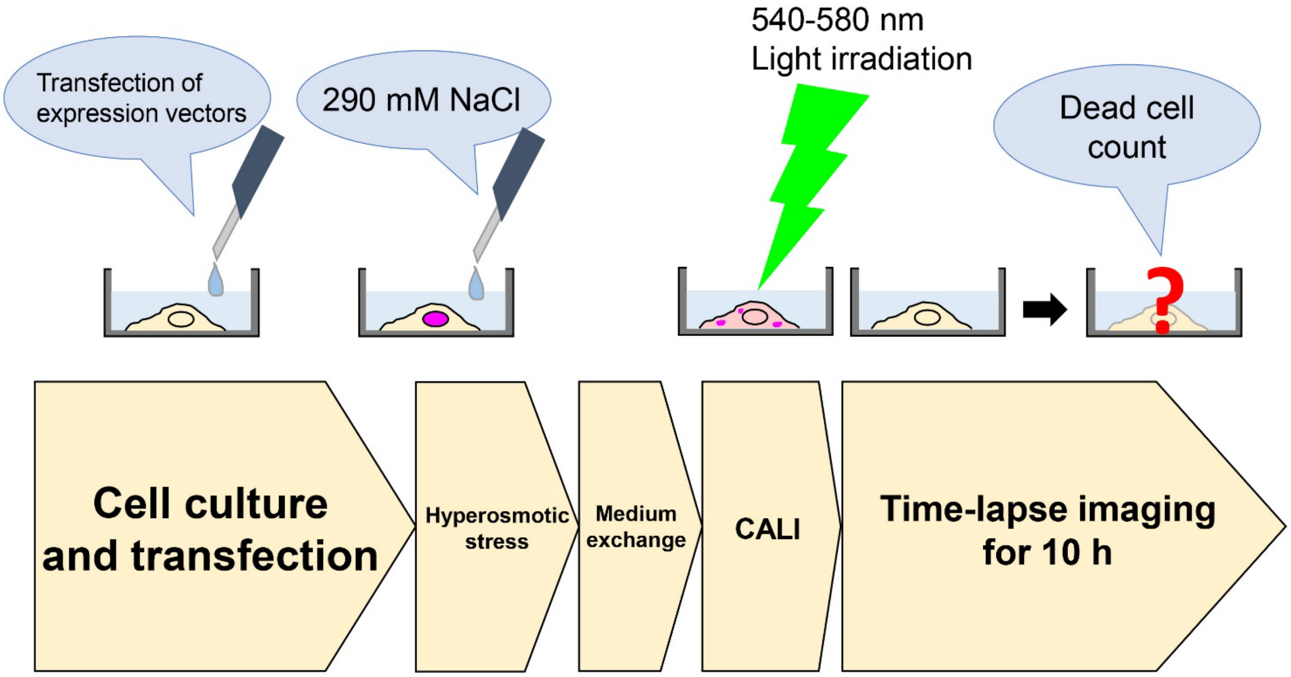


**Figure S1.** The scheme of gene transfection to the cells, hyperosmotic stress, CALI, and time-lapse observation.

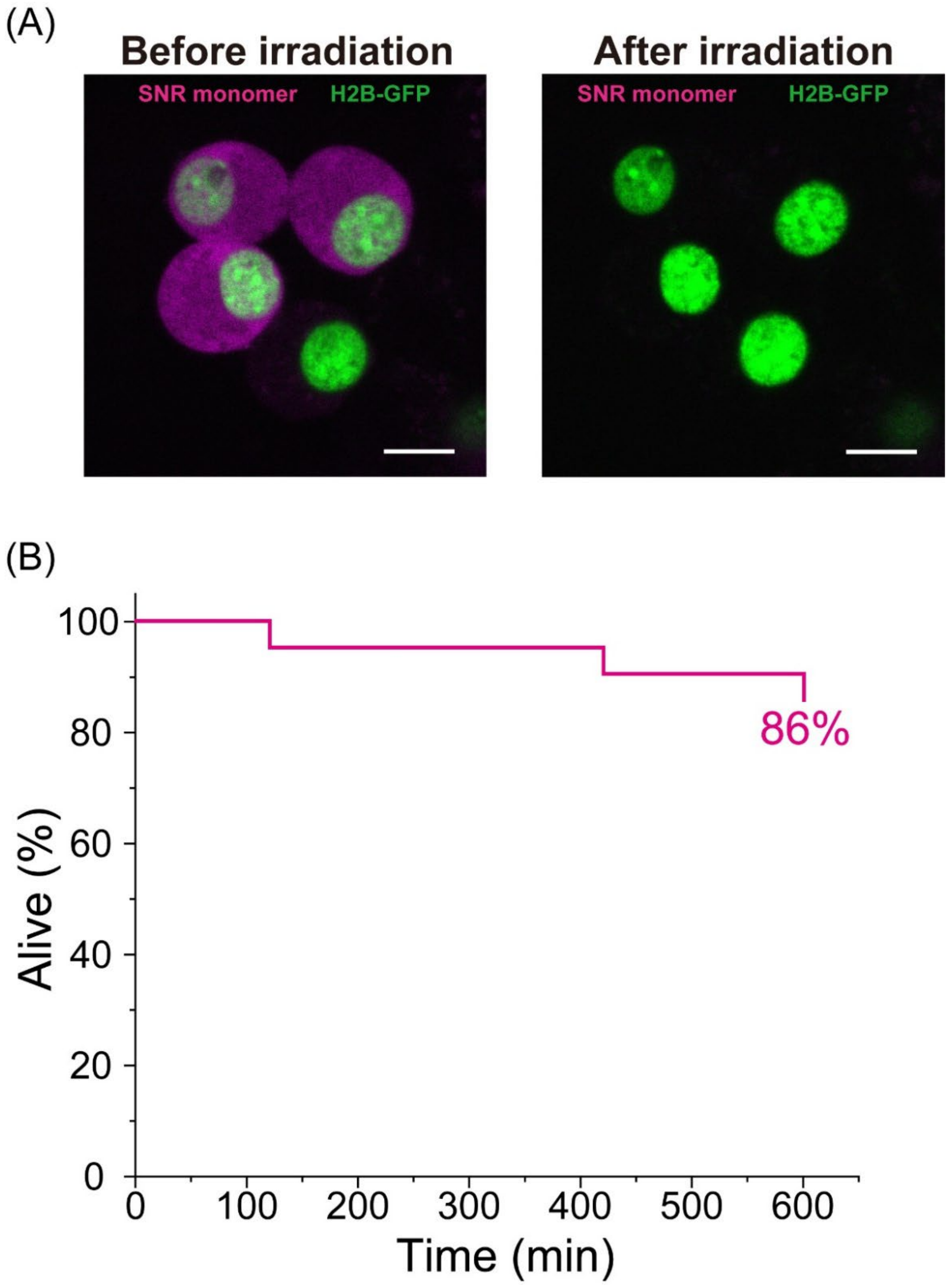


**Figure S2.** Cell viability after CALI targeting SuperNpva-Red monomers in cells in culture recovering from hyperosmotic stress. (A) Confocal fluorescent images of cells expressing SuperNova-Red monomer and Histon H2B-GFP before and after the light irradiation for CALI to SuperNova-Red. Bar = 10 µm. (B) Cell viability plot during 600 min in the recovery medium after hyperosmotic stress (21 cells). The inset number indicates the percentage of cells alive at 600 min.

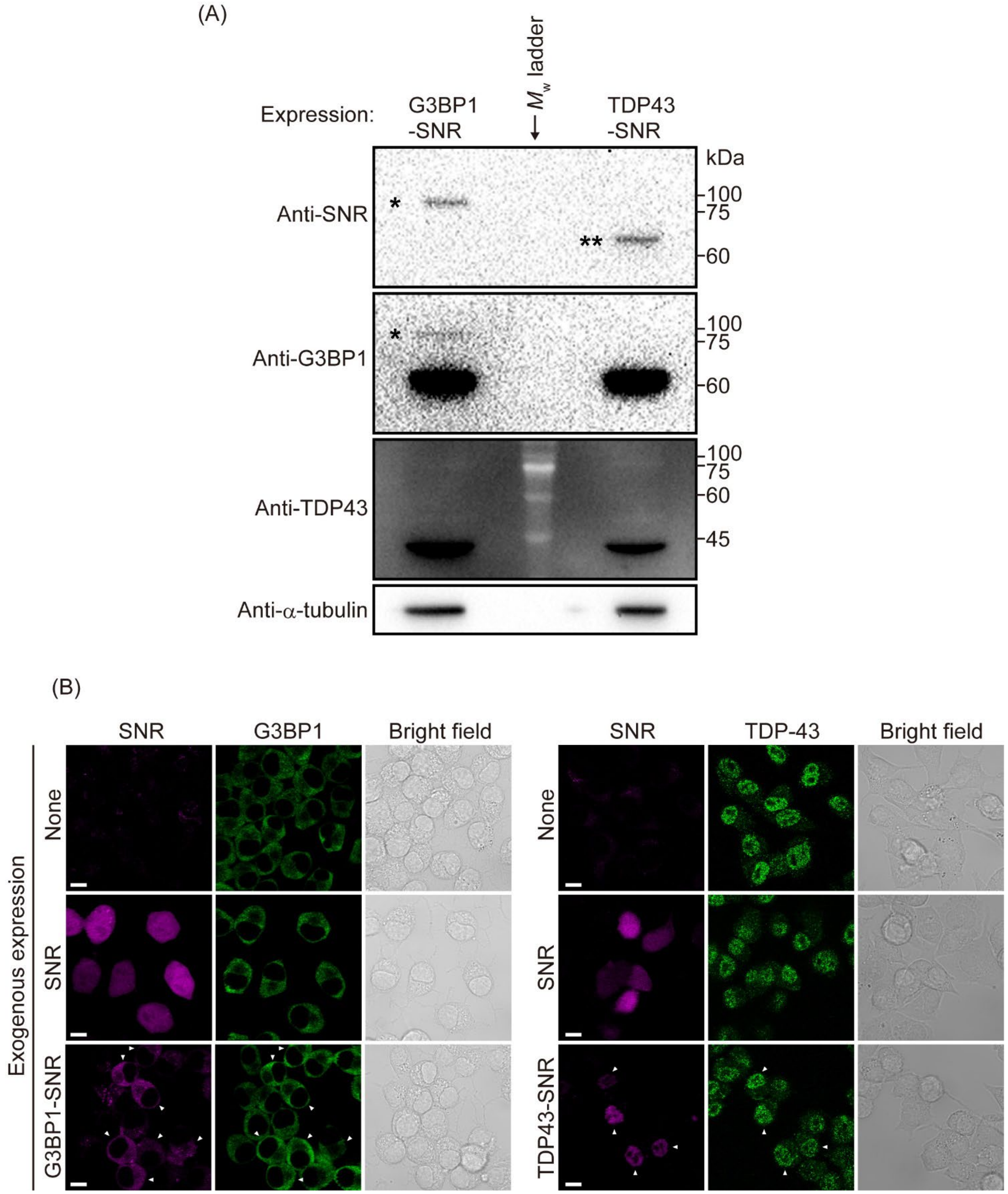


**Figure S3.** Expression level of exogenous G3BP1-SNR and TDP43-SNR in Neuro-2a cells. (A) Western blot of cell lysates expressing SuperNova-Red (SNR)-tagged G3BP1 (Left lane) or TDP-43 (Right lane) using anti-SNR, anti-G3BP1, anti-TDP-43, and anti-α-tubulin antibodies. A molecular weight marker is applied in the middle lane ($M_w$ ladder). The values on the right side of the images indicate standard molecular weights of the $M_w$ ladder. Single and double asterisks (* and **) show the observed migration of G3BP1-SNR (81 kDa) and TDP43-SNR (73 kDa), respectively. Long exposure and contrast-adjusted images were shown for G3BP1 and TDP-43 because the expression levels of SNR-tagged proteins were relatively low compared to their endogenous ones. The band of TDP43-SNR (**) was not observed in a long exposure membrane stained with an anti-TDP43 antibody due to its low expression level. (B) Immunofluorescence images of Neuro-2a cells expressing G3BP1-SNR or TDP43-SNR. SNR and Alexa Fluor 488 fluorescence was shown as magenta and green color, respectively. White arrowheads indicate the cells expressing G3BP1-SNR or TDP43-SNR. No significant increase in green fluorescence intensity of G3BP1/TDP43-SNR-positive cells was observed, indicating exogenous SNR-tagged proteins were not overexpressed. Bar = 10 μm.

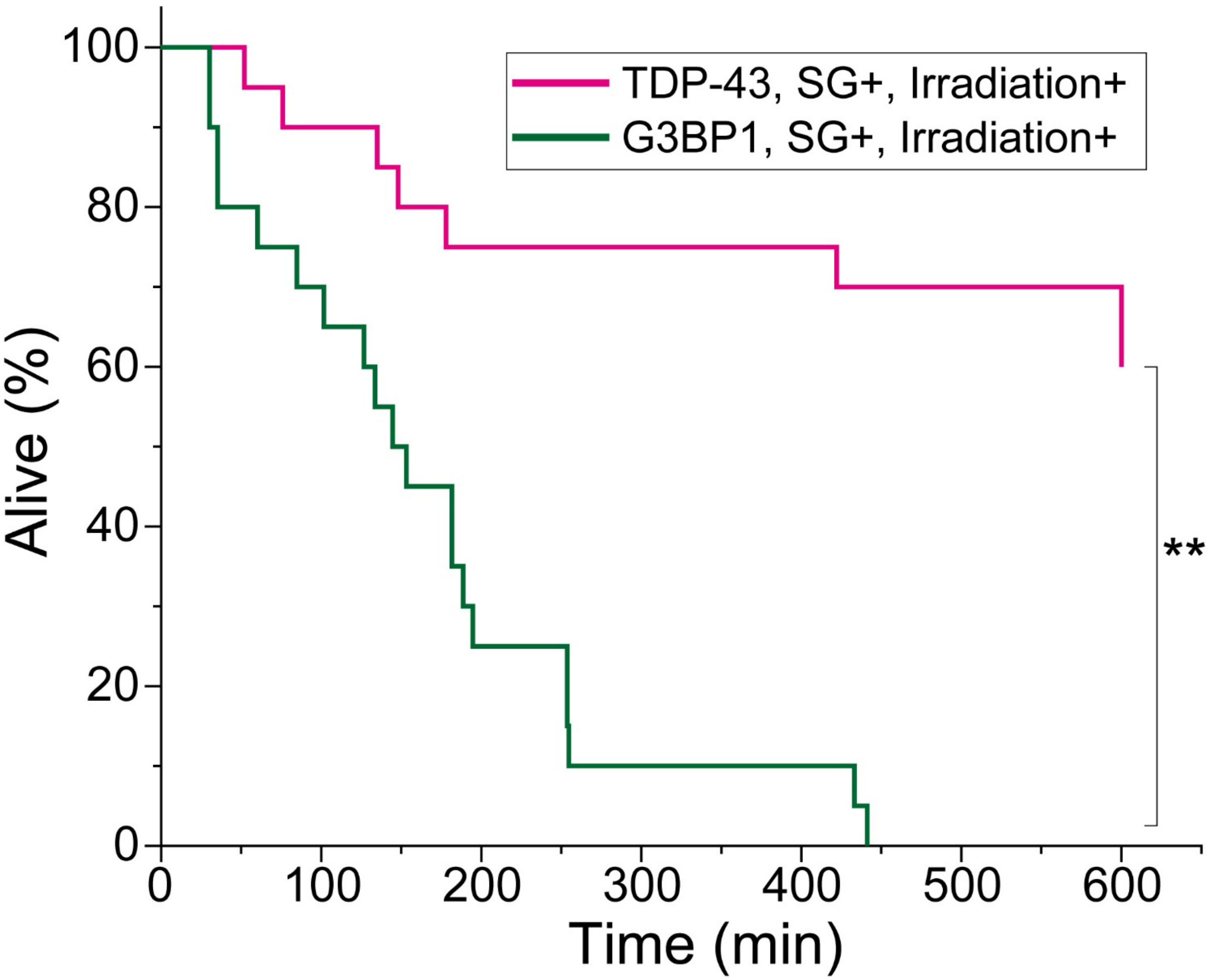


**Figure S4.** Comparison of cell viability after chromophore-associated light inactivation (CALI) between G3BP1 and TDP-43 in Neuro-2a cells recovering from hyperosmotic stress. The plot lines are derived from the data shown in Figures 1B and 2B (20 cells). P-values were obtained using the generalized Wilcoxon test (Gehan-Breslow method). **$p < 0.01$ between the lines.

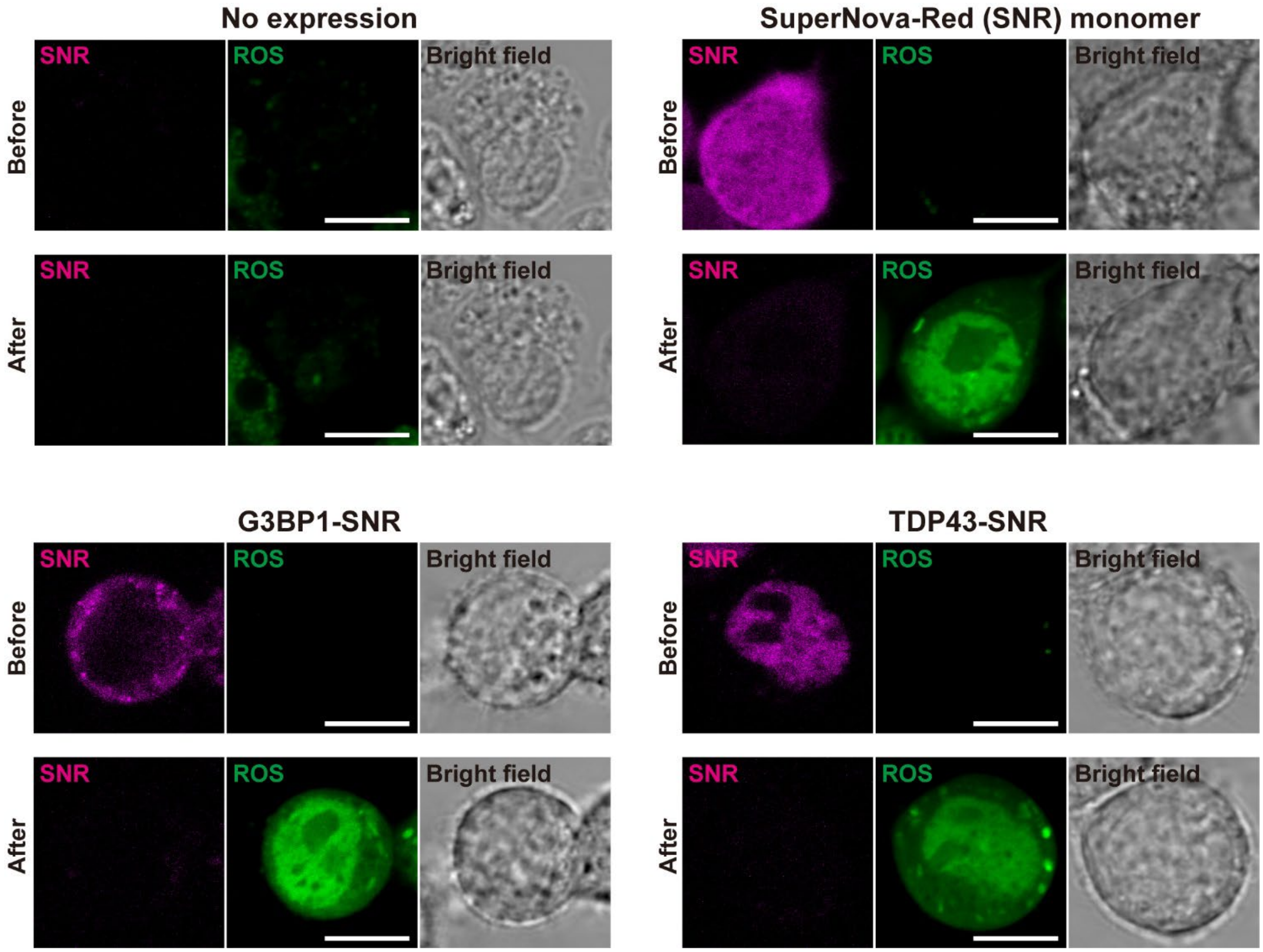


**Figure S5.** Intracellular ROS detection generated by SuperNova-Red (SNR) after light irradiation. Confocal fluorescent images of cells expressing G3BP1-SNR, TDP43-SNR, SNR monomer or nothing (no expression) before and after the light irradiation for CALI. Green fluorescence indicates ROS generation. Bar = 10 μm.

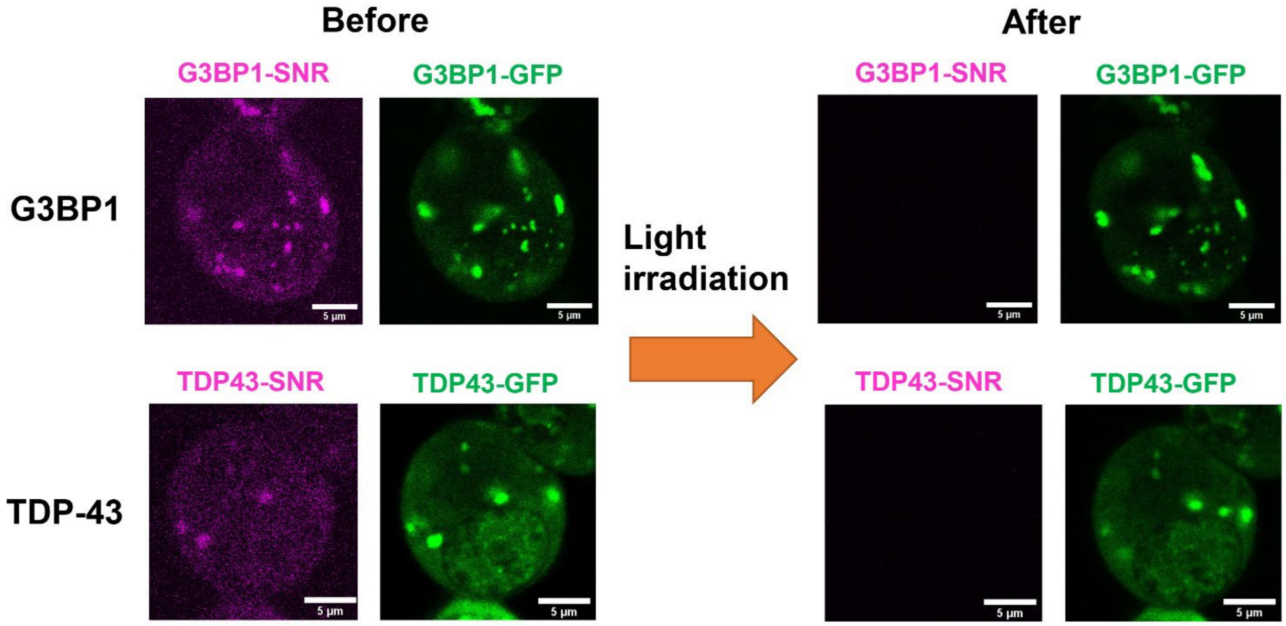


**Figure S6.** Confocal fluorescence images of cells co-expressing SuperNova-Red (SNR)- and GFP-tagged G3BP1 or TDP-43 before and after the light irradiation for CALI. Bar = 5 μm. Although slight position changes of SGs due to their three-dimensional movement were observed during the 2 minutes required for SNR photobleaching for CALI, SGs did not disappear after the light irradiation for CALI.
